# Antigenicity and infectivity characterization of SARS-CoV-2 BA.2.86

**DOI:** 10.1101/2023.09.01.555815

**Authors:** Sijie Yang, Yuanling Yu, Fanchong Jian, Weiliang Song, Ayijiang Yisimayi, Xiaosu Chen, Yanli Xu, Peng Wang, Jing Wang, Lingling Yu, Xiao Niu, Jing Wang, Tianhe Xiao, Ran An, Yao Wang, Qingqing Gu, Fei Shao, Ronghua Jin, Zhongyang Shen, Youchun Wang, Yunlong Cao

## Abstract

The recently identified SARS-CoV-2 variant, BA.2.86, which carries a substantial number of Spike mutations, has raised a global alarm. An immediate assessment of its antigenic properties and infectivity is necessary. Here, we reveal the distinct antigenicity of BA.2.86 compared with previous variants including XBB.1.5. BA.2.86 significantly evades convalescent plasma from XBB breakthrough infection (BTI) and reinfections. Key mutations that mediate the enhanced resistance include N450D, K356T, L452W, A484K, V483del, and V445H on the RBD, while BA.2.86’s NTD mutations and E554K on SD1 also largely contribute. However, we found that BA.2.86 pseudovirus exhibits compromised efficiency of infecting HEK293T-hACE2 cells compared to XBB.1.5 and EG.5, which may be caused by K356T, V483del, and E554K, and could potentially limit BA.2.86’s transmissibility. In sum, it appears that BA.2.86 has traded its infectivity for higher immune evasion during long-term host-viral evolution. Close attention should be paid to monitoring additional mutations that could improve BA.2.86’s infectivity.

## Main

The newly emerged SARS-CoV-2 saltation variant, BA.2.86, has aroused global concern (Figure 1A). Although only 24 sequences have been detected thus far, they originate from multiple countries, exhibit no epidemiological relevance, and were found in individuals without travel history, suggesting the presence of underlying international transmission (Figure 1B). On August 18th, the World Health Organization designated it as a variant under monitoring (VUM), taking into account the large number of mutations it carries ^1^. BA.2.86 harbors numerous mutations that significantly deviate from the currently designated strains, with 33 spike mutations/14 receptor binding domain (RBD) mutations relative to BA.2 and 35 spike mutations/12 RBD mutations relative to XBB.1.5 (Figure 1A and 1C). Along with the shared mutations with XBB.1.5 (T19I, 24-26del, A27S, G142D, 144del, G339H, G446S, N460K, and F486P), additional mutations I332V, K356T, V445H, N450D, N481K, A484K and 483del on BA.2.86’s RBD are likely to enhance immune evasion as previously reported ^2-6^. Many unusual mutations on the N-terminal domain (NTD), such as R21T, S50L, 69-70del, V127F, F157S, R158G, 211del, L212I, L216F, H245N, and A264D, may also alter the antigenicity of BA.2.86 ^7,8^. The above underscores the potential of BA.2.86 for global spread. Therefore, an experimental assessment of the antigenicity and infectivity of BA.2.86 is urgently needed.

**Figure 1.**
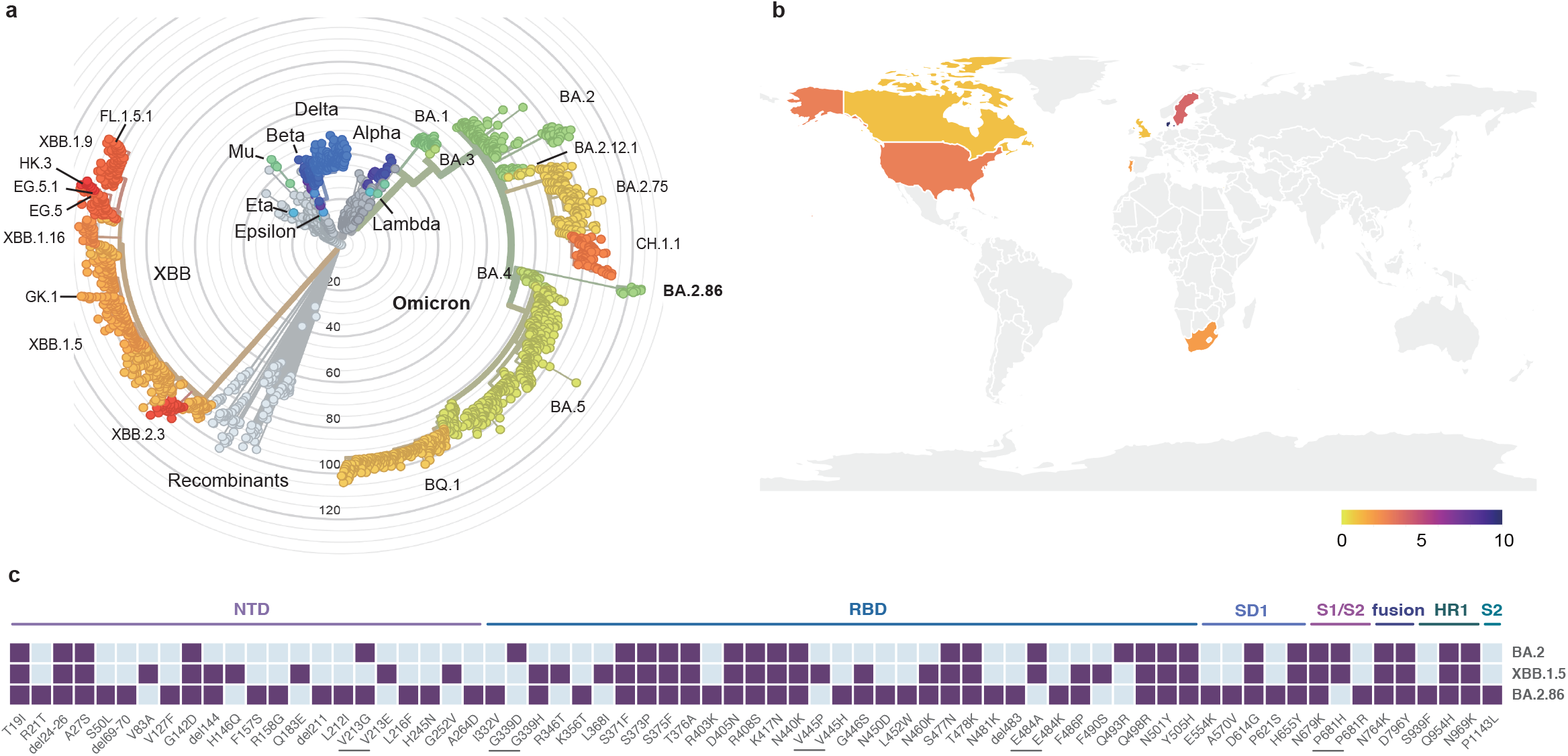
Phylogeny, prevalence and spike mutations of BA.2.86. (A) Phylogenetic tree of existing SARS-CoV-2 variants including BA.2.86. Color scales indicate the clades. Representative clades and strains are labeled. (B) Geographic distribution of detected BA.2.86 sequences. The regions where the BA.2.86 strain was not detected are denoted by gray. (C) Mutations on the spike glycoprotein of BA.2, XBB.1.5 and BA.2.86. Purple color represents the existence of mutations for each strain. The relatively absent mutations are indicated in sky blue. The spike locations of listed mutations are labeled above. The short grey lines highlight the mutations on the same sites.

First, we generated the pseudovirus of BA.2.86 and determined its antigenic distance to WT, BA.5, BQ.1.1, and XBB using serum samples from mice that had received two doses of spike mRNA vaccines (Figure 2A and S1). BA.2.86 showed high resistance to serum neutralization across all vaccine groups (Figure S1). Antigenic cartography calculated based on the pseudovirus neutralization titers showed that BA.2.86 was antigenically distinct from WT, BA.2, BA.5, and XBB.1.5, suggesting a substantial antigenic drift, which indicates that BA.2.86 could strongly evade XBB-induced antibodies (Figure 2A).

**Figure 2.**
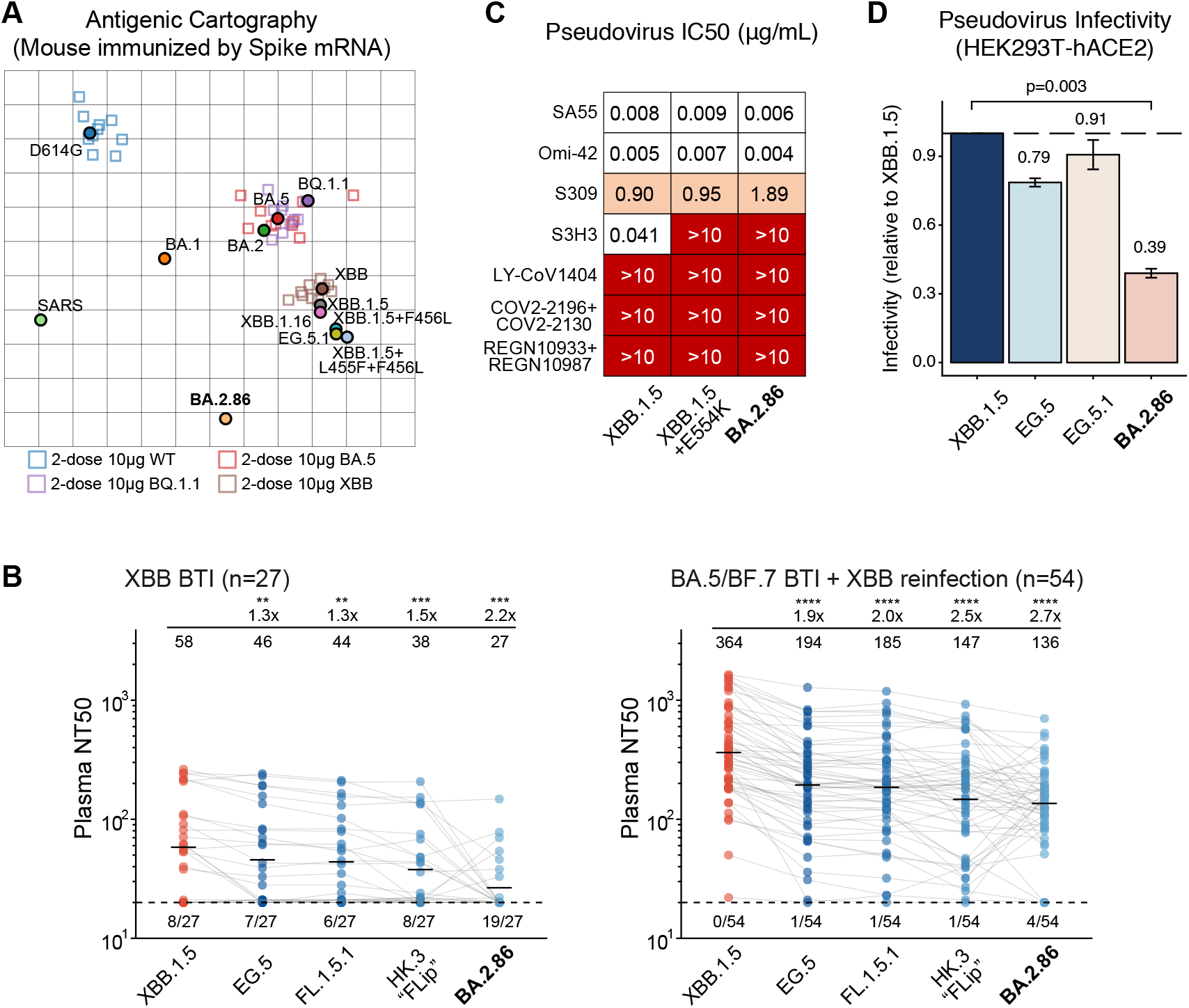
BA.2.86 exhibits distinct antigenicity and compromised infectivity. (A) Antigenic cartography built by the pseudovirus neutralization titers of mRNA-immunized mice’s plasma against various SARS-CoV-2 strains. Antigens are denoted as colored circles while plasma are shown as squares with the outlines colored by the corresponding antigens. The distances between plasma and an antigen are negatively correlated to the neutralization ability. (B) NT50 against SARS-CoV-2 XBB subvariants and BA.2.86 of convalescent plasma from individuals who received triple doses of CoronaVac and breakthrough-infected by XBB*+486P (n=27), or BA.5/BF.7 followed by XBB*+486P reinfection (n=54). Statistical significances and geometric mean titer (GMT) fold-changes are labeled in comparison with neutralization against XBB.1.5. Numbers of negative samples are labeled below the dashed line that indicates limit of detection (NT50=20). Two-tailed Wilcoxon signed-rank tests of paired samples are used. *, p<0.05; **, p<0.01; ***, p<0.001; ****, p<0.0001; NS, not significant (p>0.05). (C) IC50 (μg/mL) of approved or candidate monoclonal NAb drugs targeting RBD or SD1 on Spike, against XBB.1.5, XBB.1.5+E554K, and BA.2.86 pseudovirus. (D) Relative infectivity of EG.5, EG.5.1, and BA.2.86 compared to XBB.1.5. The efficiencies of infecting HEK293T-hACE2 cells are tested using VSV-based pseudoviruses. Error bars indicate mean±s.d. Mean values are labeled above each bar.

To assess the immune evasion characteristics of BA.2.86, pseudovirus neutralization assays are performed against XBB infection convalescent plasma and a panel of monoclonal antibodies (mAbs) (Figure 2B-C). All involved participants received three doses of inactivated vaccines before experiencing XBB (XBB*+486P) breakthrough infection (BTI). The first cohort (n=27) included individuals with single post-vaccination XBB BTI and the second cohort (n=54) comprised convalescents who experienced XBB reinfection after BA.5 or BF.7 BTI (Table S1). Indeed, we found that BA.2.86 can induce significant antibody evasion of XBB-stimulated plasma (Figure 2B). BA.2.86’s immune evasion capability even exceeds EG.5 and is comparable to “FLip” variants like HK.3 (XBB.1.5 + L455F & F456L) ^9,10^. Interestingly, the relative activities against HK.3 and BA.2.86 vary from sample to sample, indicating a large antigenic distance despite a similar level of evasion. As for monoclonal neutralizing antibody (NAb) drugs, all approved antibodies can’t neutralize BA.2.86 well, but SA55 remains effective (Figure 2C) ^11^. As expected, the E554K mutation carried by BA.2.86, which is located on the binding interface of SD1-targeting NAbs, can escape SD1-targeting NAbs, represented by S3H3, an antibody that is effective against many evasive mutants including the “FLip” variants (Figure S2A) ^12^. Importantly, we found that the E554K mutation added on XBB.1.5 could also enhance plasma evasion, suggesting that SD1-targeting NAbs compose a considerable amount in XBB-stimulated convalescent plasma (Figure S2B) ^13,14^. Furthermore, we showed that by switching the RBD part of BA.2.86 to XBB.1.5, the pseudovirus still demonstrated higher immune evasion capability than XBB.1.5 and XBB.1.5+E554K, suggesting that the mutations on BA.2.86’s NTD could also induce significant neutralizing evasion (Figure S2B). To delineate the key RBD mutations of BA.2.86’s enhanced immune evasion capability than XBB.1.5, we also tested a panel of XBB.1.5-effective NAbs against XBB.1.5-based pseudoviruses carrying single BA.2.86 RBD mutations (Figure S2C) ^15^. Results showed that N450D, K356T, L452W, A484K, V483del, and V445H are responsible for BA.2.86’s enhanced immune evasion than XBB.1.5. Specifically, K356T and L452W escape the majority of antibodies in Group E, P445H escapes D3, L452W, and A484K escape D4, and, A484K and V483del contribute to the evasion of NAbs in B/C/D1. Such systematic evasion of XBB-effective NAbs explains the distinct antigenicity of BA.2.86 and its resistance to XBB convalescent plasma. Together, the above data suggest BA.2.86 is highly immune evasive and could have advantages over currently circulating variants regarding the ability to resist XBB-induced humoral immunity.

Saltation variants may exhibit compromised efficiency of infecting host cells to gain strong capability of escaping neutralizing antibodies elicited during the antibody-virus coevolution in long-term continuous host infection ^16^. Therefore, we next evaluated BA.2.86’s cellular infectivity by testing the efficiency of its pseudovirus form in infecting hACE2-HEK293T cell lines (Figure 2D).

Surprisingly, among all tested strains, BA.2.86 exhibited the lowest infectivity compared to XBB.1.5, EG.5, and EG.5.1. Effects of the key mutations carried by BA.2.86 on infectivity relative to XBB.1.5 are also separately tested (Figure S3). As a result, the lower infectivity of BA.2.86 may be contributed mainly by K356T, V483del, and E554K. K356T introduces an N-linked glycosylation motif for N354, and V483del is near ACE2-binding sites, which may affect cell entry efficiency. Of note, the infectivity measured here is obtained through pseudovirus assays, which should be confirmed by assays using authentic BA.2.86 isolates.

In sum, we found that BA.2.86 is antigenically distinct from XBB.1.5 and previous Omicron variants, and can escape XBB-induced and XBB-effective neutralizing antibodies targeting various epitopes. Therefore, the efficacy of developing XBB-based vaccines against BA.2.86 should be closely monitored and carefully evaluated. Our results also indicate that BA.2.86 may not prevail fast due to its lower infectivity. However, BA.2.86 may obtain additional mutations during its transmission to enhance the infectivity and become predominant in the near future, just like the previous convergent evolution of S486P in XBB subvariants, which highlights the necessity of global cooperation to track the evolution of BA.2.86 ^17^.

## Acknowledgments

We are grateful to scientists in the community for their continuous tracking of SARS-CoV-2 variants and helpful discussion, including Daniele Focosi, Ryan Hisner, Cornelius Roemer, Federico Gueli, Tom Peacock, Raj Rajnarayanan, and many other researchers. We thank all volunteers for providing blood samples. This project is financially supported by the Ministry of Science and Technology of China (2023YFC3041500;2023YFC3043200), Changping Laboratory (2021A0201; 2021D0102), and the National Natural Science Foundation of China (32222030).

## Author Contributions

Y.C. designed and supervised the study. S.Y., F.J., and Y.C. wrote the manuscript with inputs from all authors. Q.G. proofread the manuscript. S.Y., W.S. and F.J. performed sequence analysis and illustration. Y.Y. and Youchun W. constructed pseudoviruses. P.W., L.Y., T.X., R.A., Yao W., J.W. (BIOPIC), J.W. (Changping Laboratory) and F.S. processed the plasma samples and performed the pseudovirus neutralization assays. W.S., A.Y., X.N., and Y.C. analyzed the neutralization data. Y.X., X.C., Z.S., and R.J. recruited the SARS-CoV-2 convalescents.

## Methods

### Plasma isolation

Blood samples were acquired from individuals who had recovered from SARS-CoV-2 Omicron BTI or experienced reinfection. These samples were collected following approved study protocols from Beijing Ditan Hospital, Capital Medical University (Ethics Committee archiving No. LL-2021-024-02), the Tianjin Municipal Health Commission, and the Ethics Committee of Tianjin First Central Hospital (Ethics Committee archiving No. 2022N045KY). All participants provided written informed consent, allowing for the collection, storage, and utilization of blood samples for research and data publication purposes. For patients in the reinfection groups, their initial infections occurred in December 2022 in Beijing and Tianjin during the BA.5/BF.7 wave[5]. From the sequences obtained between 12/01/2022 and 02/01/2023, over 98% were identified as BA.5* (excluding BQ*). The prevalent subtypes in China during that time were BA.5.2.48* and BF.7.14*, which lacked additional mutations on the RBD region, making them generally representative of BA.5/BF.7 variants. Patients in the XBB BTI cohort and those experiencing second infections in the reinfection groups encountered infections between May and June 2023, a period when over 90% of sequenced samples from Beijing and Tianjin corresponded to XBB* + 486P variants. The confirmation of these infections was carried out using PCR tests or antigen tests.

To process the blood samples, whole blood was mixed at a 1:1 ratio with PBS+2% FBS and then subjected to Ficoll gradient centrifugation (Cytiva). Following centrifugation, plasma was extracted from the upper layer. These plasma samples were divided into aliquots, stored at temperatures of - 20 °C or lower, and underwent heat inactivation prior to experimental procedures.

### Infectivity Assay

Following quantification using RT-PCR, the pseudotyped virus was diluted to achieve an equal particle count. Subsequently, 100 mL aliquots of the diluted virus were introduced into individual wells of 96-well cell culture plates. Cells from the specific cell lines being tested were then enzymatically detached using trypsin and added to each well at a concentration of 2.3 x 104 cells per 100 ml. Chemiluminescence monitoring was conducted following a 24-hour incubation period with a CO2 concentration of 5% and a temperature of 37°C. The supernatant for each sample was adjusted to a volume of 100 mL to ensure consistency. A mixture of luciferase substrate and cell lysis buffer (Perkinelmer, Fremont, CA) was prepared and added to each well at a volume of 100 mL. 150 mL of the resulting lysate was transferred to opaque 96-well plates after 2 minutes. PerkinElmer Ensight luminometer was used to detect the luminescence signal, and the data was recorded in terms of relative luminescence unit (RLU) values. Each experimental group consisted of two replicates, and the entire set of experiments was repeated three times to ensure the reliability and consistency of results.

### Antigenic cartography construction

Antigenic cartography, or antigenic map, is a well-established method utilized for quantifying and visualizing neutralization data [6]. Briefly, after preprocessing and normalization of the neutralization titer data, modified multidimensional scaling (MDS) is performed to arrange the antigen and plasma points within a single map. To construct the antigenic cartography, we used the pseudovirus NT50 titers of mRNA-immunized mice’s plasma against various strains, which include D614G, BA.1, BA.2, BA.5, BQ.1.1, XBB, XBB.1.16, XBB.1.5, EG.5.1, XBB.1.5+F456L, XBB.1.5+L455F+F456L, BA.2.86 and SARS. The mice were immunized with 2 doses (10μg) spike mRNA of WT, BA.5 BQ.1.1 or XBB, and serum was sampled one week after the last immunization. Antigenic cartography was computed by the R package Racmacs(v1.1.35), and visualized by the R package ggplot2(v3.4.1). The construction of the maps involved 500 optimization runs, with the minimum column basis parameter set to “none.”

### Neutralization assays

A vesicular stomatitis virus (VSV) pseudovirus packaging system was employed to generate Spike pseudovirus variants of SARS-CoV-2. The spike plasmid of each variant was integrated into the pcDNA3.1 vector. The G*ΔG-VSV virus (VSV G pseudotyped virus, obtained from Kerafast) and the spike protein plasmid were introduced into 293T cells (American Type Culture Collection [ATCC], CRL-3216) through transfection. Once cultured, the resulting pseudovirus was collected from the supernatant, subjected to filtration, divided into aliquots, and then frozen at a temperature of −80°C for later utilization.

For pseudovirus neutralization assays, the Huh-7 cell line (Japanese Collection of Research Bioresources [JCRB], 0403) was employed. Plasma samples or antibodies were subjected to serial dilution in the culture media and combined with the pseudovirus. The mixture was then incubated at 37°C with 5% CO_2_ for 1 hour. Subsequently, digested Huh-7 cells were introduced into the antibody-virus mixture. Following a day of incubation within the culture chamber, the supernatant was discarded. D-luciferin reagent (PerkinElmer, 6066769) was added to the plates and allowed to incubate in darkness for 2 minutes. Cell lysis content was then transferred to detection plates. The luminescence intensity was measured using a microplate spectrophotometer (PerkinElmer, HH3400). The IC50 value was calculated using a four-parameter logistic regression model.

## Supplementary Tables and Figures

**Table S1** | **Information of SARS-CoV-2 convalescent patients involved in the study**.

**Figure S1.**
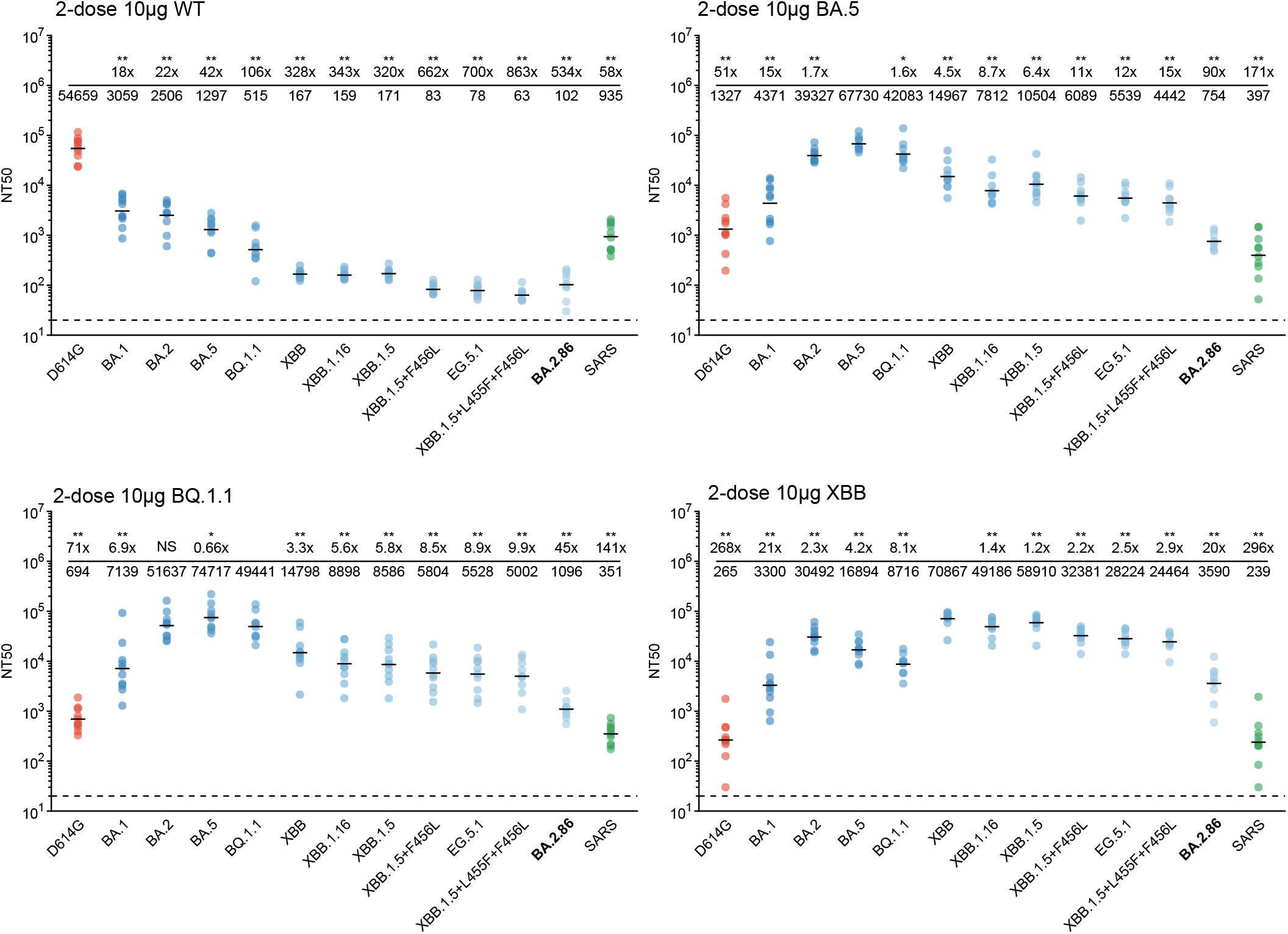
Neutralization titers of immunized mice sera. NT50 of sera from mice immunized by SARS-CoV-2 WT, BA.5, BQ.1.1, or XBB Spike mRNA, against SARS-CoV-2 variants. Data are used to construct the antigenic cartography in Figure 2B. Statistical significances and geometric mean titer (GMT) fold-changes are labeled in comparison with neutralization against XBB.1.5. Two-tailed Wilcoxon signed-rank tests of paired samples are used. *, p<0.05; **, p<0.01; NS, not significant (p>0.05).

**Figure S2.**
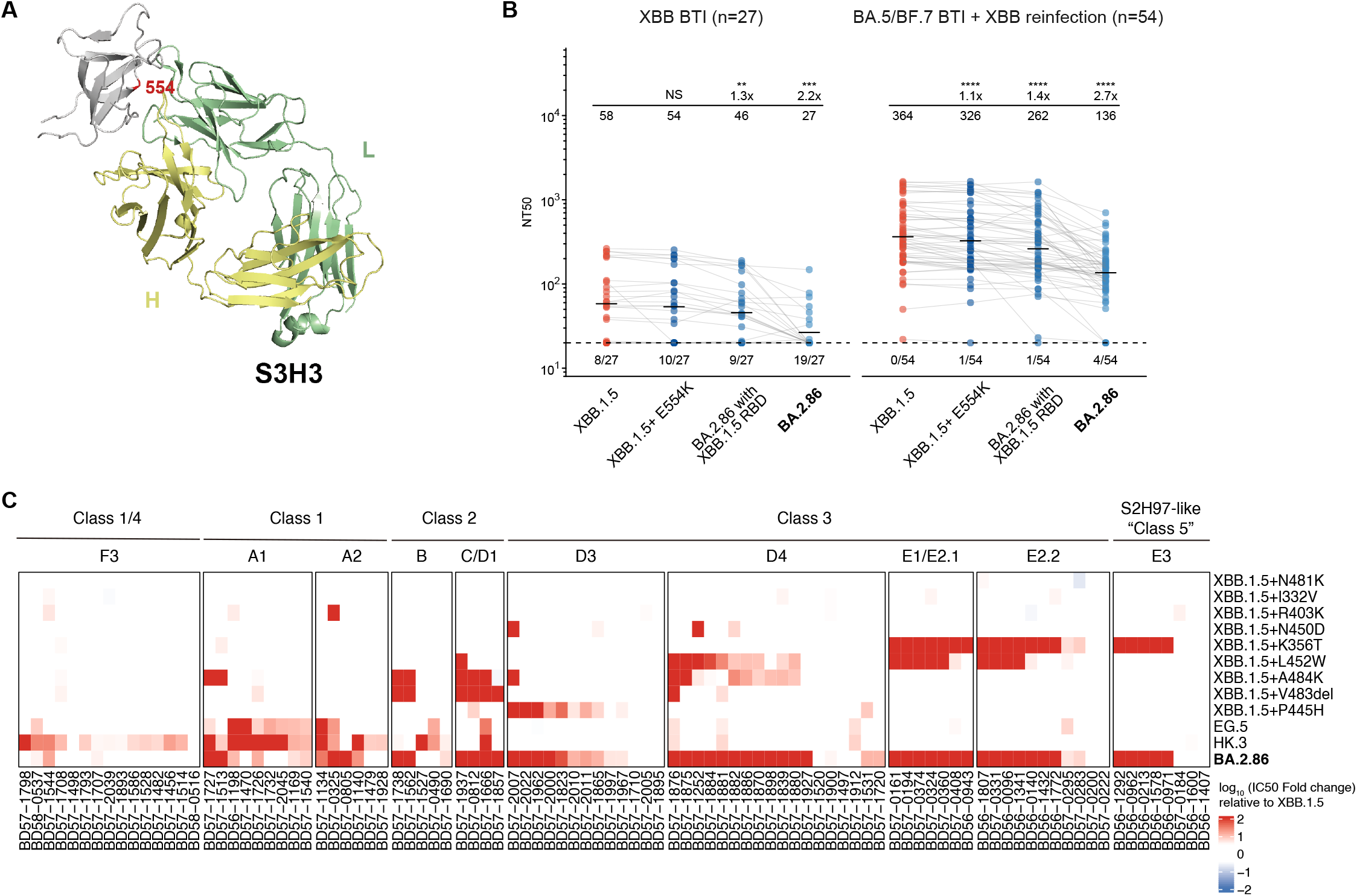
Distinct antigenicity of BA.2.86. (A) Structure of SD1-targeting antibody S3H3 in complex with Beta Spike (PDB: 7WDF). E554, which is mutated to Lys in BA.2.86, is on the interface between S3H3 and SD1, leading to escape. (B) NT50 against XBB.1.5, XBB.1.5+E554K, and BA.2.86 with XBB.1.5 RBD of convalescent plasma from individuals who received triple doses of CoronaVac and breakthrough-infected by XBB*+486P (n=27), or BA.5/BF.7 followed by XBB*+486P reinfection (n=54). Statistical significances and geometric mean titer (GMT) fold-changes are labeled in comparison with neutralization against XBB.1.5. Numbers of negative samples are labeled below the dashed line that indicates limit of detection (NT50=20). Two-tailed Wilcoxon signed-rank tests of paired samples are used. *, p<0.05; **, p<0.01; ***, p<0.001; ****, p<0.0001; NS, not significant (p>0.05). Neutralization against BA.2.86 (Figure 2B) is shown here again for comparison. (C) Pseudovirus neutralization of a panel of XBB.1.5-effective monoclonal NAbs targeting different RBD epitopes (determined by DMS) against XBB subvariants and XBB.1.5 with BA.2.86 mutations. Fold changes compared to IC50 against XBB.1.5 are shown as a heatmap with epitope groups and classes labeled above.

**Figure S3.**
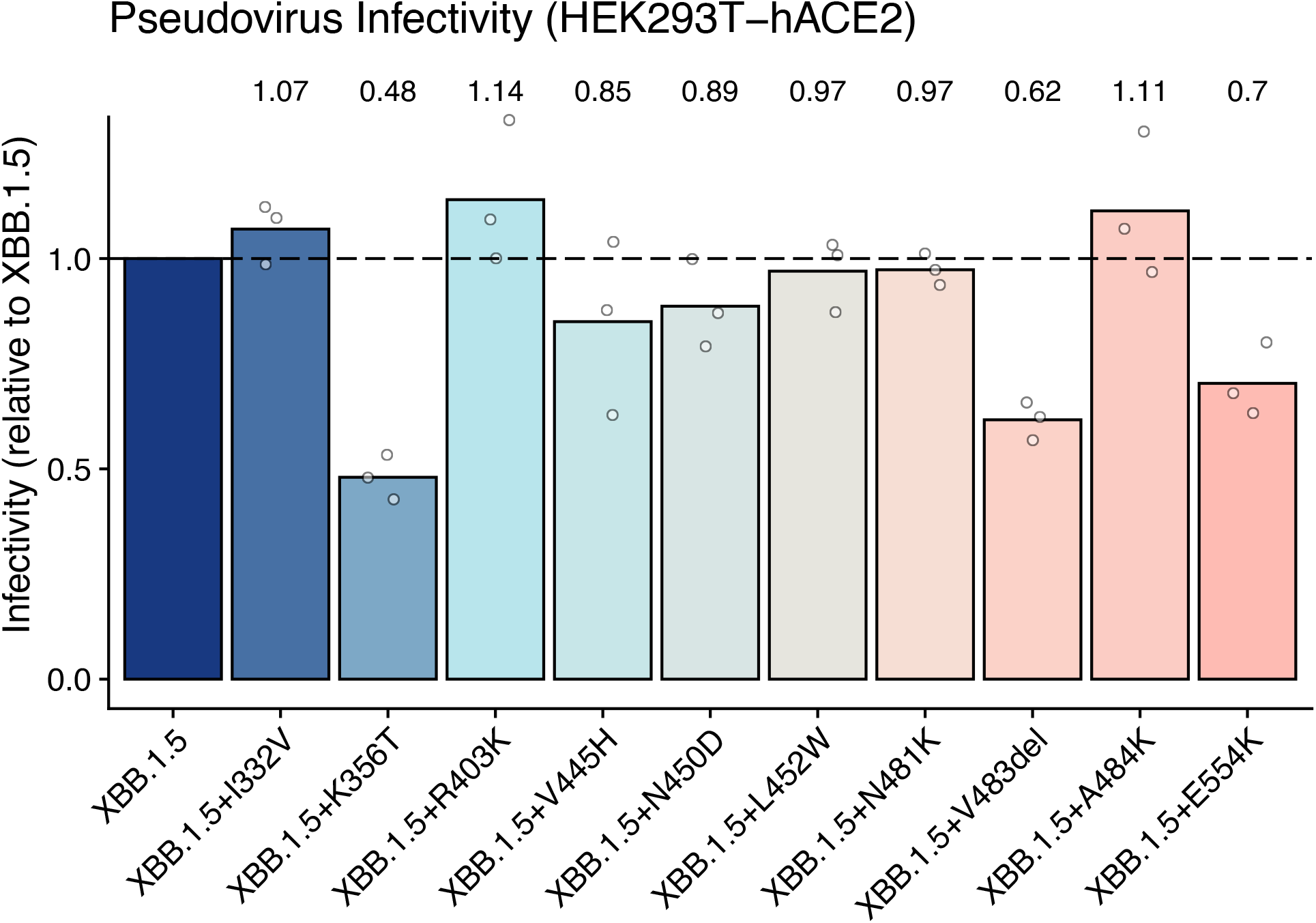
Relative infectivity of XBB.1.5 with BA.2.86 mutations. Relative infectivity of XBB.1.5 pseudovirus with mutations carried by BA.2.86. The efficiencies of infecting HEK293T-hACE2 cells compared to that of XBB.1.5 are shown. Error bars indicate mean±s.d. Mean values are labeled above each bar. Experiments were conducted in three biological replicates.

## References

1. News, U. (2023). WHO, CDC Tracking New COVID-19 Variant BA.2.86. https://www.usnews.com/news/health-news/articles/2023-08-18/who-cdc-tracking-new-covid-19-variant-ba-2-86.

2. Greaney, A.J., Starr, T.N., Barnes, C.O., Weisblum, Y., Schmidt, F., Caskey, M., Gaebler, C., Cho, A., Agudelo, M., Finkin, S., et al. (2021). Mapping mutations to the SARS-CoV-2 RBD that escape binding by different classes of antibodies. Nature Communications 12, 4196. 10.1038/s41467-021-24435-8.

3. Cao, Y., Wang, J., Jian, F., Xiao, T., Song, W., Yisimayi, A., Huang, W., Li, Q., Wang, P., An, R., et al. (2022). Omicron escapes the majority of existing SARS-CoV-2 neutralizing antibodies. Nature 602, 657–663. 10.1038/s41586-021-04385-3.

4. Cao, Y., Yisimayi, A., Jian, F., Song, W., Xiao, T., Wang, L., Du, S., Wang, J., Li, Q., Chen, X., et al. (2022). BA.2.12.1, BA.4 and BA.5 escape antibodies elicited by Omicron infection. Nature 608, 593–602. 10.1038/s41586-022-04980-y.

5. Greaney, A.J., Starr, T.N., and Bloom, J.D. (2022). An antibody-escape estimator for mutations to the SARS-CoV-2 receptor-binding domain. Virus Evol 8, veac021. 10.1093/ve/veac021.

6. Ragonnet-Cronin, M., Nutalai, R., Huo, J., Dijokaite-Guraliuc, A., Das, R., Tuekprakhon, A., Supasa, P., Liu, C., Selvaraj, M., Groves, N., et al. (2023). Generation of SARS-CoV-2 escape mutations by monoclonal antibody therapy. Nature Communications 14, 3334. 10.1038/s41467-023-37826-w.

7. Cao, Y., Song, W., Wang, L., Liu, P., Yue, C., Jian, F., Yu, Y., Yisimayi, A., Wang, P., Wang, Y., et al. (2022). Characterization of the enhanced infectivity and antibody evasion of Omicron BA.2.75. Cell Host Microbe. 10.1016/j.chom.2022.09.018.

8. Wang, Q., Iketani, S., Li, Z., Guo, Y., Yeh, A.Y., Liu, M., Yu, J., Sheng, Z., Huang, Y., Liu, L., and Ho, D.D. (2022). Antigenic characterization of the SARS-CoV-2 Omicron subvariant BA.2.75. Cell Host Microbe, 2022.2007.2031.502235. 10.1016/j.chom.2022.09.002.

9. Jian, F., Yang, S., Yu, Y., Song, W., Yisimayi, A., Chen, X., Xu, Y., Wang, P., Yu, L., Wang, J., et al. (2023). Convergent evolution of SARS-CoV-2 XBB lineages on receptor-binding domain 455-456 enhances antibody evasion and ACE2 binding. bioRxiv, 2023.2008.2030.555211. 10.1101/2023.08.30.555211.

10. Kaku, Y., Kosugi, Y., Uriu, K., Ito, J., Kuramochi, J., Sadamasu, K., Yoshimura, K., Asakura, H., Nagashima, M., Consortium, T.G.t.P.J., and Sato, K. (2023). Antiviral efficacy of the SARS-CoV-2 XBB breakthrough infection sera against Omicron subvariants including EG.5. bioRxiv, 2023.2008.2008.552415. 10.1101/2023.08.08.552415.

11. Cao, Y., Jian, F., Zhang, Z., Yisimayi, A., Hao, X., Bao, L., Yuan, F., Yu, Y., Du, S., Wang, J., et al. (2022). Rational identification of potent and broad sarbecovirus-neutralizing antibody cocktails from SARS convalescents. Cell Rep 41, 111845. 10.1016/j.celrep.2022.111845.

12. Hong, Q., Han, W., Li, J., Xu, S., Wang, Y., Xu, C., Li, Z., Wang, Y., Zhang, C., Huang, Z., and Cong, Y. (2022). Molecular basis of receptor binding and antibody neutralization of Omicron. Nature 604, 546–552. 10.1038/s41586-022-04581-9.

13. Bianchini, F., Crivelli, V., Abernathy, M.E., Guerra, C., Palus, M., Muri, J., Marcotte, H., Piralla, A., Pedotti, M., De Gasparo, R., et al. (2023). Human neutralizing antibodies to cold linear epitopes and subdomain 1 of the SARS-CoV-2 spike glycoprotein. Science Immunology 8, eade0958. doi:10.1126/sciimmunol.ade0958.

14. Seow, J., Khan, H., Rosa, A., Calvaresi, V., Graham, C., Pickering, S., Pye, V.E., Cronin, N.B., Huettner, I., Malim, M.H., et al. (2022). A neutralizing epitope on the SD1 domain of SARS-CoV-2 spike targeted following infection and vaccination. Cell Reports 40, 111276. 10.1016/j.celrep.2022.111276.

15. Yisimayi, A., Song, W., Wang, J., Jian, F., Yu, Y., Chen, X., Xu, Y., Yang, S., Niu, X., Xiao, T., et al. (2023). Repeated Omicron exposures override ancestral SARS-CoV-2 immune imprinting. bioRxiv, 2023.2005.2001.538516. 10.1101/2023.05.01.538516.

16. Kemp, S.A., Collier, D.A., Datir, R.P., Ferreira, I.A.T.M., Gayed, S., Jahun, A., Hosmillo, M., Rees-Spear, C., Mlcochova, P., Lumb, I.U., et al. (2021). SARS-CoV-2 evolution during treatment of chronic infection. Nature 592, 277–282. 10.1038/s41586-021-03291-y.

17. Yue, C., Song, W., Wang, L., Jian, F., Chen, X., Gao, F., Shen, Z., Wang, Y., Wang, X., and Cao, Y. (2023). ACE2 binding and antibody evasion in enhanced transmissibility of XBB.1.5. The Lancet Infectious Diseases 23, 278–280. 10.1016/S1473-3099(23)00010-5.

